# Entner–Doudoroff pathway and Non-OxPPP bypasses glycolysis and OxPPP in *Ralstonia solanacearum*

**DOI:** 10.1101/2020.01.31.929778

**Authors:** Poonam Jyoti, Manu Shree, Chandrakant Joshi, Tulika Prakash, Suvendra Kumar Ray, Siddhartha Sankar Satapathy, Shyam Kumar Masakapalli

## Abstract

In *Ralstonia solanacearum*, a devastating phytopathogen whose metabolism is poorly understood, we observed that Entner-Doudoroff (ED) pathway and NonOxidative pentose phosphate pathway (OxPPP) bypasses glycolysis and OxPPP under glucose oxidation. Evidences derived from ^13^C stable isotopes feeding and genome annotation based comparative metabolic network analysis supported the observations. Comparative metabolic network analysis derived from the currently available *53* annotated *R. solanacearum* strains also including the recently reported strain (F1C1), representing the four phylotypes confirmed the lack of key genes coding for phosphofructokinase (*pfk-1*) and phosphogluconate dehydrogenase (*gnd*) enzymes that are relevant for glycolysis and OxPPP respectively. *R. solanacearum* F1C1 cells fed with ^13^C Glucose (99%[1-^13^C]- or 99%[1,2-^13^C]- or 40%[^13^C_6_]-glucose) followed by GC-MS based labelling analysis of fragments from amino acids, glycerol and ribose provided clear evidence that rather than Glycolysis and OxPPP, ED pathway and NonOxPPP are the main routes sustaining metabolism in *R. solanacearum*. The ^13^C incorporation in the mass ions of alanine (m/z 260, m/z 232); valine (m/z 288, m/z 260), glycine (m/z 218), serine (m/z 390, m/z 362), histidine (m/z 440, m/z 412), tyrosine (m/z 466, m/z 438), phenylalanine (m/z 336, m/z 308), glycerol (m/z 377) and ribose (m/z 160) mapped the pathways supporting the observations. The outcomes help better defining the central carbon metabolic network of *R. solanacearum* that can be integrated with ^13^C metabolic flux analysis as well as flux balance analysis studies for defining the metabolic phenotypes.

**Importance:** Understanding the metabolic versatility of *Ralstonia solanacearum* is important as it regulates the tradeoff between virulence and metabolism (1, 2) in a wide range of plant hosts. Due to a lack of clear evidence until this work, several published research papers reported on potential roles of Glycolysis and Oxidative pentose phosphate pathways (OxPPP) in *R. solanacearum* (3, 4). This work provided evidence from ^13^C stable isotopes feeding and genome annotation based comparative metabolic network analysis that Entner-Doudoroff pathway and Non-OxPPP bypasses glycolysis and OxPPP during the oxidation of Glucose, one of the host xylem pool that serves as a potential carbon source (5). The outcomes help better defining the central carbon metabolic network of *R. solanacearum* that can be integrated with ^13^C metabolic flux analysis as well as flux balance analysis studies for defining the metabolic phenotypes. The study highlights the need to critically examine phytopathogens whose metabolism is poorly understood.

## Introduction

*Ralstonia solanacearum* is one of the most destructive plant pathogen as it infects over 450 plant species (6–8) and its metabolism is poorly understood. It is a soil borne pathogen that enters the plants through natural openings or wounds, colonize and blocks water conduction in the xylem which leads to wilting and death of plants (9). Recent studies showed that there is a resource allocation or tradeoff between metabolism and virulence (1–2). Carbon metabolism not only play significant role in bacterial growth but also involves in extracellular polysaccharide production and hrp gene expression which are crucial for virulence and pathogenicity (10, 11). While the systems features of virulence are widely studied, no detailed investigations on the central metabolism of *R. solanacearum* is undertook. Owing to its wide host range wherein the pathogen encounters varying nutritional regimes, it is very important to understand its metabolic features. Metabolic versatility of the bacteria plays crucial role in conferring growth as well as pathogenicity in its complex life cycle. This is supported by studies on the differential expression of genes in pathogenic and non-pathogenic strain of *R. solanacearum* wherein about 50% of the genes belong to carbohydrate and amino acid metabolism (9-14). Annotations based on genome analysis and stable isotope labelling can map the central metabolic pathways of *R. solanacearum*. Mapping of the oxidation of different carbon sources via the central metabolic pathways will shed light on how the cellular demands of NADPH, ATP and other cofactors are met via metabolism eventually defining the metabolic phenotypes.

The rich repository of available genomes of different *R. solanacearum* phylotypes or strains obtained via the Next-generation sequencing (NGS) can readily be annotated and comparatively analyzed using bioinformatics to decode the metabolic features. Currently, the genome data of 127 *R. solanacearum* strains is available in the database of National Center for Biotechnology Information (NCBI) out of which 47 are complete genome sequences, 13 chromosome sequences, 32 scaffolds and 35 contigs (NCBI, as on Oct 2019 https://www.ncbi.nlm.nih.gov/genome/genomes/490). Also 10 *R. solanacerum* annotated genomes are available in the databases KEGG (Kyoto encyclopedia of genes and genomes)(15) and Microscope/Genoscope(16). In this study, 53 complete genome sequences of *R. solanacearum* covering phylotypes (I, II III and IV) were selected for comparative pathway analysis. In addition, 10 other *Ralstonia spp*. strains were included for analysis. Studies have shown that the *R. solanacearum* is of around 5-6 MB with a chromosome and a megaplasmid that houses various genes related to virulence, metabolism, regulatory and secretory systems.

In this work, comparative genome analysis has predicted the absence of some key pathway genes and enzymes (see results section), whose validations of the pathway activities were further derived from ^13^C tracer-based mapping studies. Given the complexity of carbon transitions in the networks, often parallel ^13^C feeding experiments would be needed to get a clear understanding of the pathways as well as in generating experimental evidence to confirm the missing genes if any. For ^13^C based retro biosynthetic mapping of metabolic pathway activities, Gas Chromatography Mass

Spectroscopy (GC-MS) based isotopomer analysis of different aminoacid fragments derived from protein hydrolysates would be needed (17). Experimental plan, selection of stable isotopes and measuring the isotope labelling in carbon atoms of the metabolites are crucial steps for pathway mapping using ^13^C approach (18-19).

Since Glucose is one of the preferred carbon sources available in host xylem pool for *R. solanacearum* metabolism (2, 5), we adopted the comparative genome and ^13^C mapping studies to validate and map central metabolic pathways of glycolysis, Oxidative Pentose phosphate pathway (OxPPP), Non-oxidative pentose phosphate pathway (Non-OxPPP) and Entner-Doudoroff (ED) pathway. We subjected *R. solanacearum* cells to [1- ^13^C]-, [1,2- ^13^C]- and [^13^C_6_]-glucose and investigated the label incorporation in the amino acids to gain comprehensive insight into the glucose oxidation pathways of central metabolism. The study generated comprehensive evidence to conclude that Entner-Doudoroff pathway and Non-OxPPP bypasses glycolysis and OxPPP in *Ralstonia solanacearum*.

## Results

### Comparative pathway analysis of *Ralstonia solanacearum* strains highlighted the absence of key central metabolic pathway genes

Central metabolic pathways (glycolysis, Entner-Doudoroff (ED) pathway and pentose phosphate pathway) of the selected organisms were reconstructed and visualised based on KEGG Orthology (KO) identifiers using KEGG pathway reconstruct tool. The analysis confirmed the completeness of ED and Non-OxPPP reactions (Figure 1, Supplementary Table S1). It was observed that all the selected *Ralstonia spp* (except *R. picketti*) lacked *pfk* gene that codes for an important regulatory enzyme phosphofructokinase-1 (EC 2.7.1.11, KO0850), indicating the possible absence of glucose oxidation by glycolytic pathway. Also, *gnd* gene coding for phosphogluconate dehydrogenase enzymes (EC 1.1.1.44 and EC 1.1.1.343, KO0033) is absent in all *Ralstonia spp* studied indicating incomplete OxPPP (Supplementary Table S1). Experimental evidence in relation to the absence of these pathways in *R. solanacearum* is generated by parallel feeding of multiple [^13^C]glucose substrates and tracking the label redistribution.

**Figure 1.**
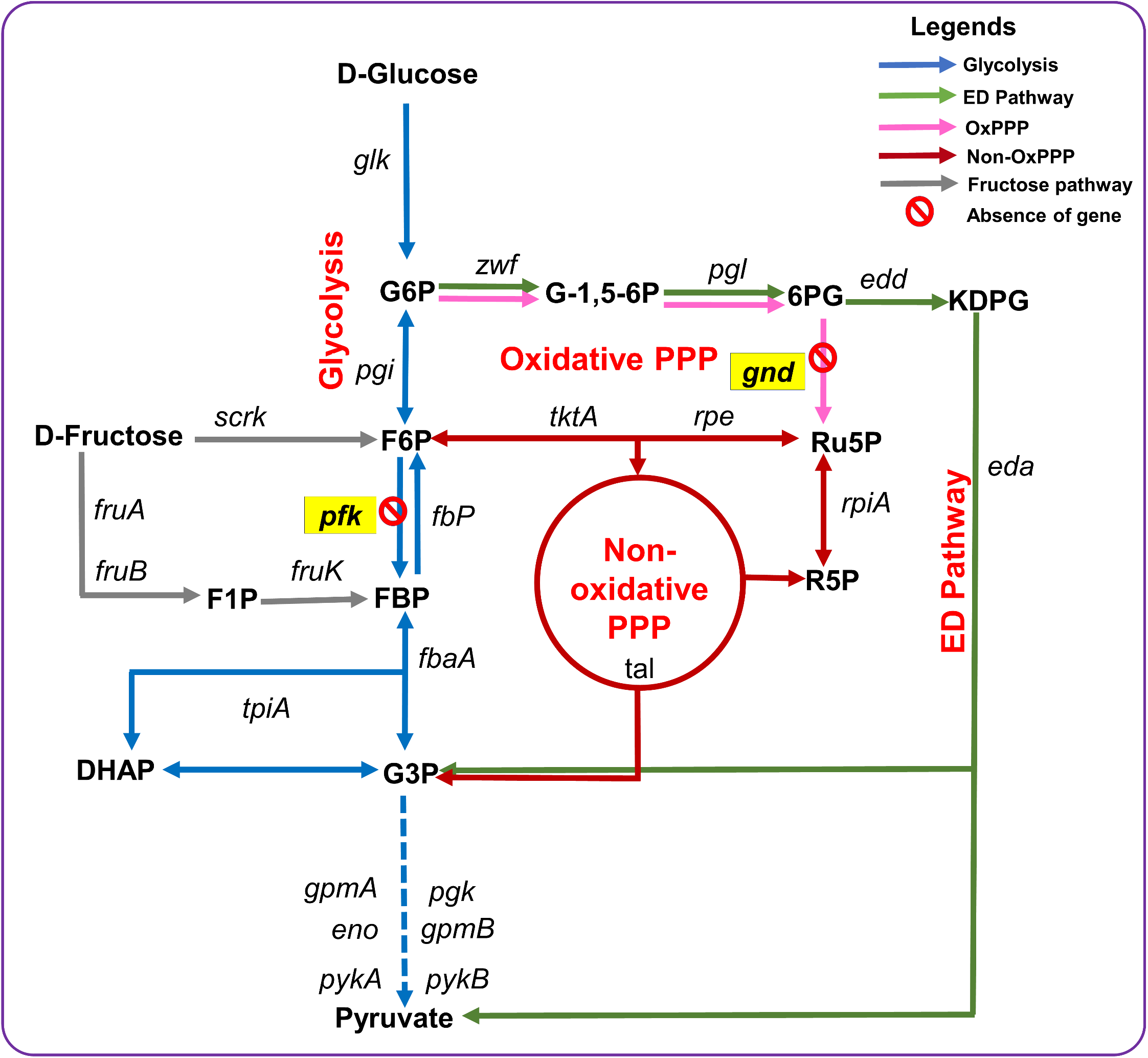
Glucose oxidation pathways in *Ralstonia solanacearum* strains highlight absence of key pathway genes. Central metabolic pathways mainly glycolysis, Entner-Doudoroff (ED) and pentose phosphate (PP) pathway among 63 strains of *Ralstonia spp*. were mapped for the annotation of relevant genes based on comparative genome analysis (see Supplementary Table S1). All *Ralstonia solanacearum* strains (53) lacked glycolysis and Oxidative Pentose Phosphate pathway (OxPPP) genes *pfk and gnd* that codes for phosphofructokinase and 6PG dehydrogenase respectively. Among all the 63 strains of *Ralstonia spp*. studied only *R. picketti* (see Supplementary Table S1) showed presence of *pfk*. The missing genes are highlighted in yellow colored boxes (*gnd, pfk*). Reactions between metabolites in the pathways, namely, glycolysis, ED pathway, OxPPP, Non-OxPPP and fructose metabolic pathway are shown in blue, green, pink, brown and grey colored arrows respectively. Genes coding the respective enzymes to catalyze the reactions and metabolites are presented in abbreviated form, The abbreviation used are: G6P: glucose-6-phosphate; F6P: fructose-6-phosphate; FBP: fructose 1,6 bisphosphate; G3P: glyceraldehyde 3-phosphate; DHAP: Dihydroxyacetone phosphate; G-1,5-6P: Glucono-1-5-lactone-6phosphate; 6PG: phosphogluconate; KDPG: 2-keto-3-deoxy-6phosphogluconate; Ru5P: Ribulose 5Phosphate; R5P: Ribose 5phosphate; *glk:* glucokinase; *pgi*: glucose-6-phosphate isomerase; *pfk:* phosphofructokinase; *tpi*: triosephosphate isomerase; *pgk*: phosphoglycerate kinase; *gpm*: phosphoglycerate mutase; *eno*: enolase; *pyk*: pyruvate kinase; *zwf*: G6P dehydrogenase; *pgl*: phospho gluconolactonoase; *edd*: phosphogluconate dehydratase; *gnd*: 6PG dehydrogenase; *eda*: 2-dehydro-3-deoxyphosphogluconate aldolase, *tkt*: transketolase; *rpe*: ribulose-phosphate 3-epimerase; *rpi*: ribose 5-phosphate isomerase; *tal*: transaldolase; *fbp*: fructose-1,6-bisphosphataseI; *fruK*: 1-phosphofructokinase; *fruA, fruB*: phosphotransferase system, enzyme I, *scrk*: fructokinase and *fba*A: fructose bisphosphate aldolase, class II.

### Amino acid fragments reporting on central metabolic pathways identified

The TBDMS derivatised protein hydrolysate of [^12^C]-, [1^−13^C]-, [1,2^−13^C]- and 40%[^13^C_6_]- glucose fed cells were subjected to GC-MS for accurate analysis of mass isotopomer distribution in amino acids. The total ion chromatogram of protein hydrolysate resulted in 15 amino acids (Figure 2). Each TBDMS-derivatised amino acid due to ionization formed different fragments in the mass spectroscopy such as [M-0]^+^, [M-15]^+^, [M-57]^+^, [M-85]^+^, [M-159]^+^ and [M-R]^+^ (R denotes the side chain of an amino acid often resulting in fragment [f302]^+^). The 15 amino acids with their respective elution time (min), the reliable mass ions (m/z) of fragments [M-85] and [M-57] along with the carbon numbers are presented in Supplementary Table S2.

**Figure 2.**
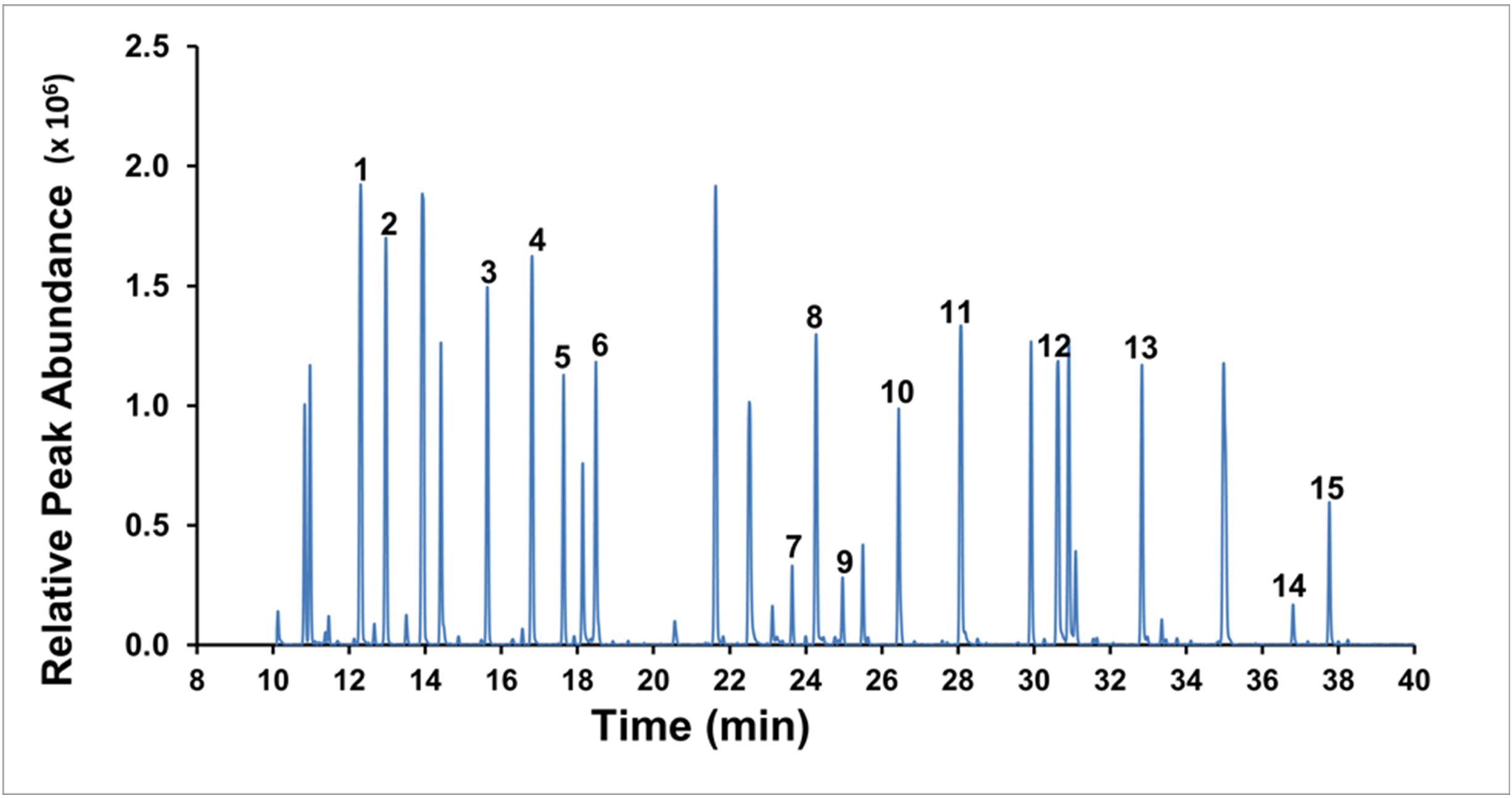
Total ion chromatogram of TBDMS derivatized protein hydrolysate of *R. solanacearum* obtained by GC-MS. The peaks corresponding to 15 aminoacids are numbered. The detected aminoacids with their respective elution time are: Alanine (12.285); Glycine (12.958), Valine (15.622), leucine (16.793), Isoleucine (17.629), proline (18.476), Methionine (23.632), Serine (24.262), Threonine (24.955), Phenylalanine (26.436), Aspartic acid (28.068), Glutamic acid (30.612), Lysine (32.826), Histidine (36.793) and Tyrosine (37.752). The retention time, derivative, specific ions and carbon backbone of each aminoacid fragment detected is presented in Supplementary Table S2.

We observed that [M-57] fragment ion contains all the backbone carbons of an amino acid, while first carbon is missing in [M-85] fragment. The fragments that are valid for further analysis were derived based on average ^13^C analysis (Supplementary Table S3). The average ^13^C abundances (%) of each amino acid fragment as well as relative mass isotopomer distributions (MIDs) used for pathway mapping is presented in Supplementary Tables S3 and S4 respectively. The ^13^C feeding studies has shown substantial isotope label incorporation in the amino acids of *R. solanacearum*. In cells fed with 40% [^13^C_6_]glucose, most of the amino acids except glycine resulted in average ^13^C enrichment to the extent of 35%. In case of glycine the average ^13^C labelling was ~20% highlighting the potential dilution from either external CO_2_ or Methylenetetrahydrofolate due to glycine synthase reaction, where glycine cleave to CO_2_, NH^4+^, and a methylene group (–CH2–), which is accepted by tetrahydrofolate (THF) in a reversible reaction (20). The mass isotopomers of amino acids derived from positional [1^−13^C]-, [1,2^−13^C]- glucose fed *R. solanacearum* under same conditions as that of [^13^C_6_]glucose supported comprehensive analysis.

### ED and Non-OxPPP bypasses glycolysis and OxPPP

Average ^13^C incorporation in amino acid fragments retro-biosynthetically reported on central metabolite precursors thereby highlighting the glucose oxidation pathway activities in *R. solanacearum* mainly via ED and Non-OxPPP (Figure 3, Figure 4 and Figure 5, Supplementary Table S3 & S4). Mainly the label incorporation in [M-57] and/or [M-85] fragments of Alanine m/z 260, m/z 232; Valine m/z 288, m/z 260; Glycine m/z 218; Serine m/z 390, m/z 362; Histidine m/z 440 m/z 412; Tyrosine m/z 466, m/z 438; Phenylalanine m/z 336, m/z 308; glycerol m/z 377, ribose m/z 160 provided the relevant evidence.

**Figure 3.**
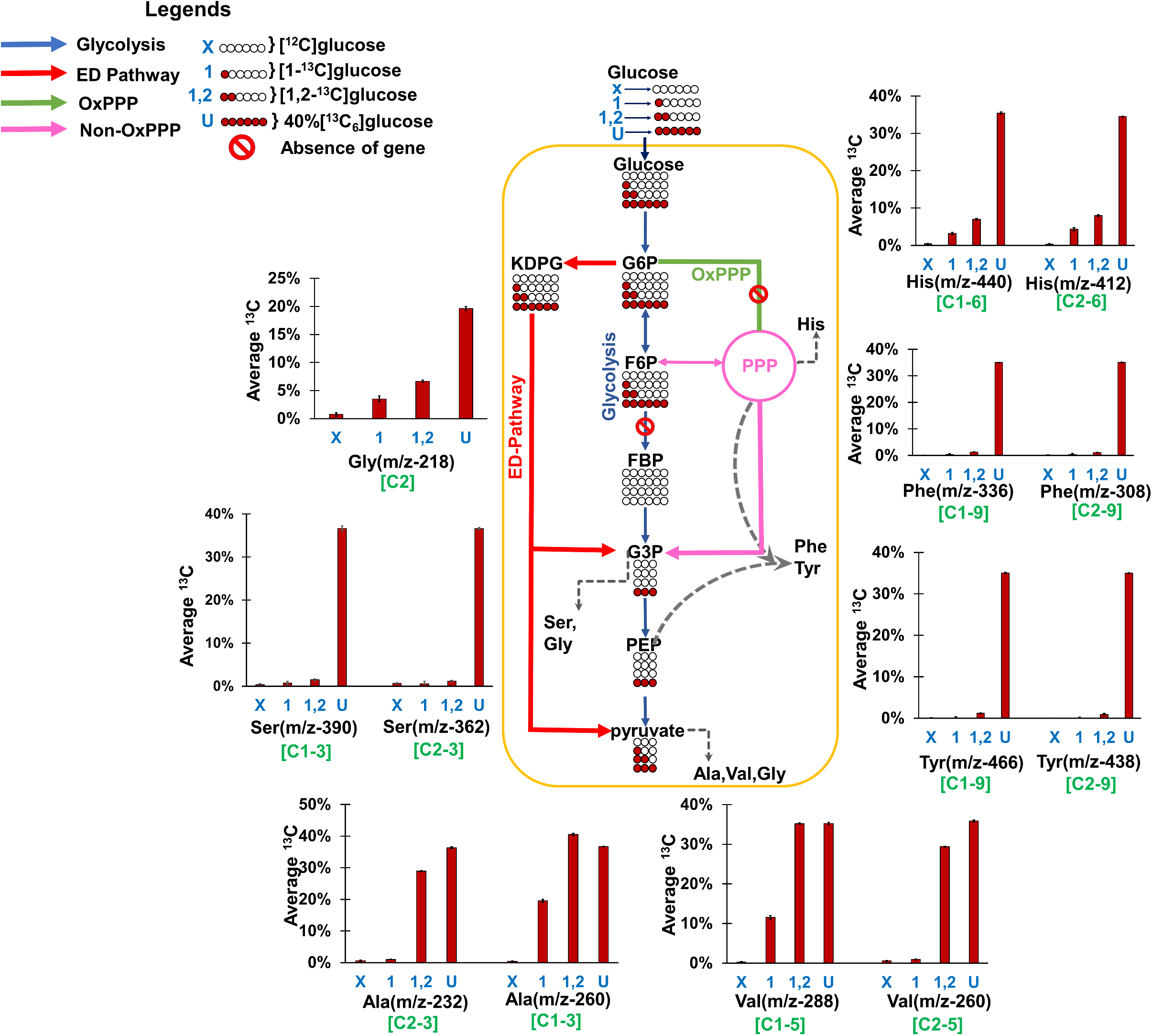
Average ^13^C label incorporation in the amino acids of *R. solanacearum* F1C1. The abbreviations “X”, “1”, “1,2” and “U” represents the cells fed with [^12^C] glucose (X), [1-^13^C] glucose (1), [1,2-^13^C] glucose (1,2), 40%[^13^C_6_] glucose (U) respectively. The levels of average %^13^C in amino acids ([M-57] and/or [M-85] fragments from GC-MS) highlight the extent of label incorporation which further retro biosynthetically report on each metabolite precursors as well as pathway activities. In case of X (i.e [^12^C]glucose), the average %^13^C in all fragments were at natural abundance (~1.13%). The average ^13^C in alanine, valine, serine and glycine from all parallel feeding experiments confirm the activity of ED pathway which bypasses glycolysis (see results section for detailed explanation). The circles below the metabolites corresponds to the carbon backbone with each position either labelled (red filled) or unlabelled (white) corresponding to a feeding experiment (X; 1; 1,2 or U). The reactions, namely glycolysis, ED (Entner-Doudoroff) pathway, oxidative pentose phosphate pathway (OxPPP), reductive pentose phosphate pathway and amino acids are presented in blue, red, green, pink and dotted grey color respectively. The abbreviations used for the metabolites are as follows: G6P: glucose-6-phosphate; F6P: fructose-6-phosphate; FBP: fructose 1,6 bisphosphate; G3P: glyceraldehyde 3-phosphate; PEP: phosphoenolpyruvate; KDPG: 2-keto-3-deoxy-6phosphogluconate; Ala: alanine; Val: valine; Ser: serine; Gly: glycine; Tyr: tyrosine; Phe: phenylalanine; His: Histidine.

**Figure 4.**
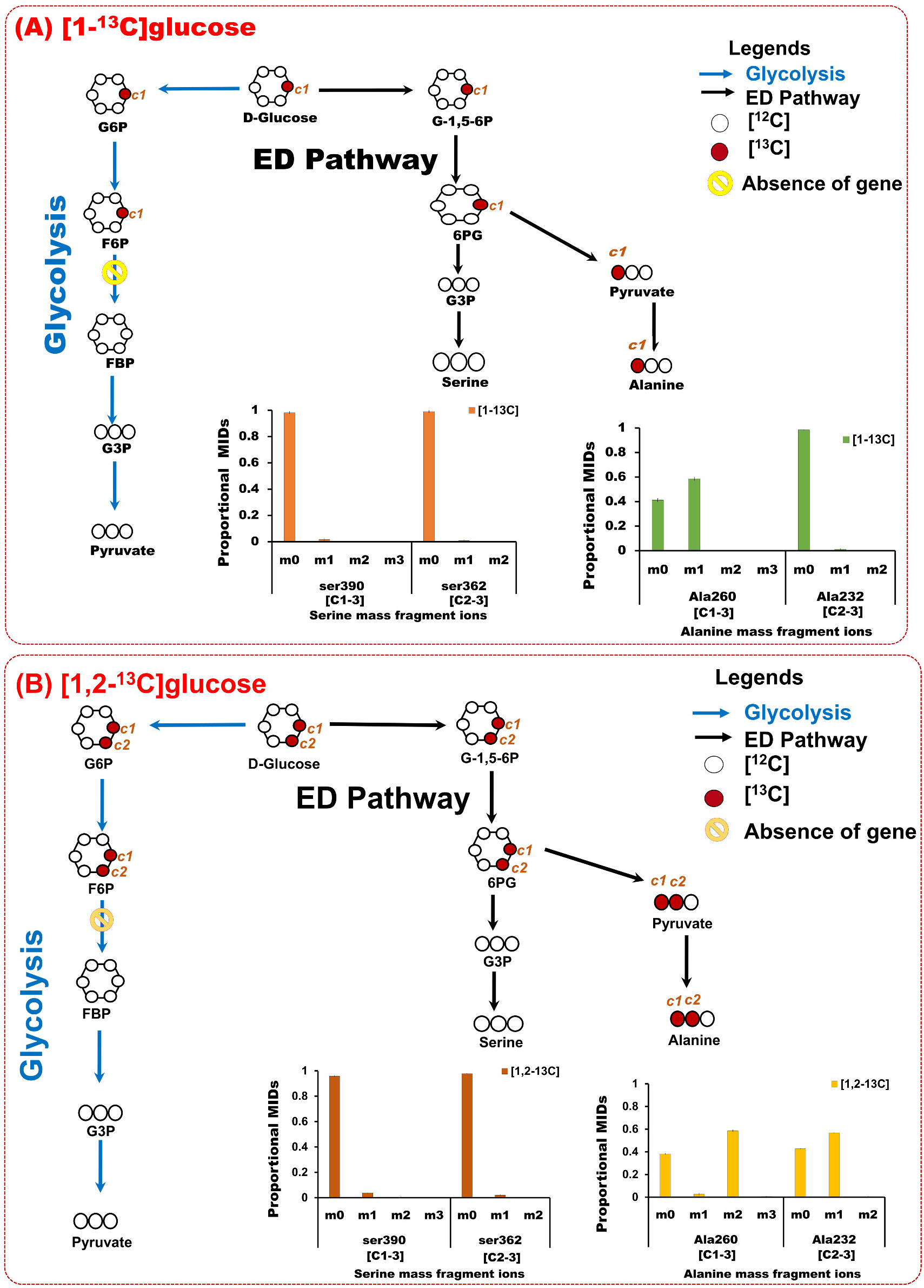
Tracking of ^13^C label in serine and alanine through glycolysis and ED pathway in *R. solanacearum* F1C1. The label enrichment in the pathway intermediates interpreted from the mass isotopomers of serine and alanine derived from cells parallelly fed with [1-^13^C]glucose (A) and [1,2-^13^C]glucose (B) are presented. The mass isotopomer distribution in serine m/z 390 (C 1-3), m/z 362 (C 2-3) and alanine m/z 260 (C1-3), m/z 232 (C 2-3) shows ED pathway is the predominant route of glucose oxidation with no appreciable activity of glycolysis ([1-^13^C]glucose oxidation via glycosis would result in labelled serine whereas ED pathway would result in unlabelled serine which is the case. Cells fed with [1,2-^13^C]glucose would also result in similar labelling pattern. Abreviations: G-1,5-6P: Glucono-1-5-lactone-6phosphate; 6PG: phosphogluconate; G6P: Glucose 6 phosphate; F6P: Fructose 6 phosphate; G3P: Glyceraldehyde 3 phosphate; MID: mass isotopomer distribution

**Figure 5.**
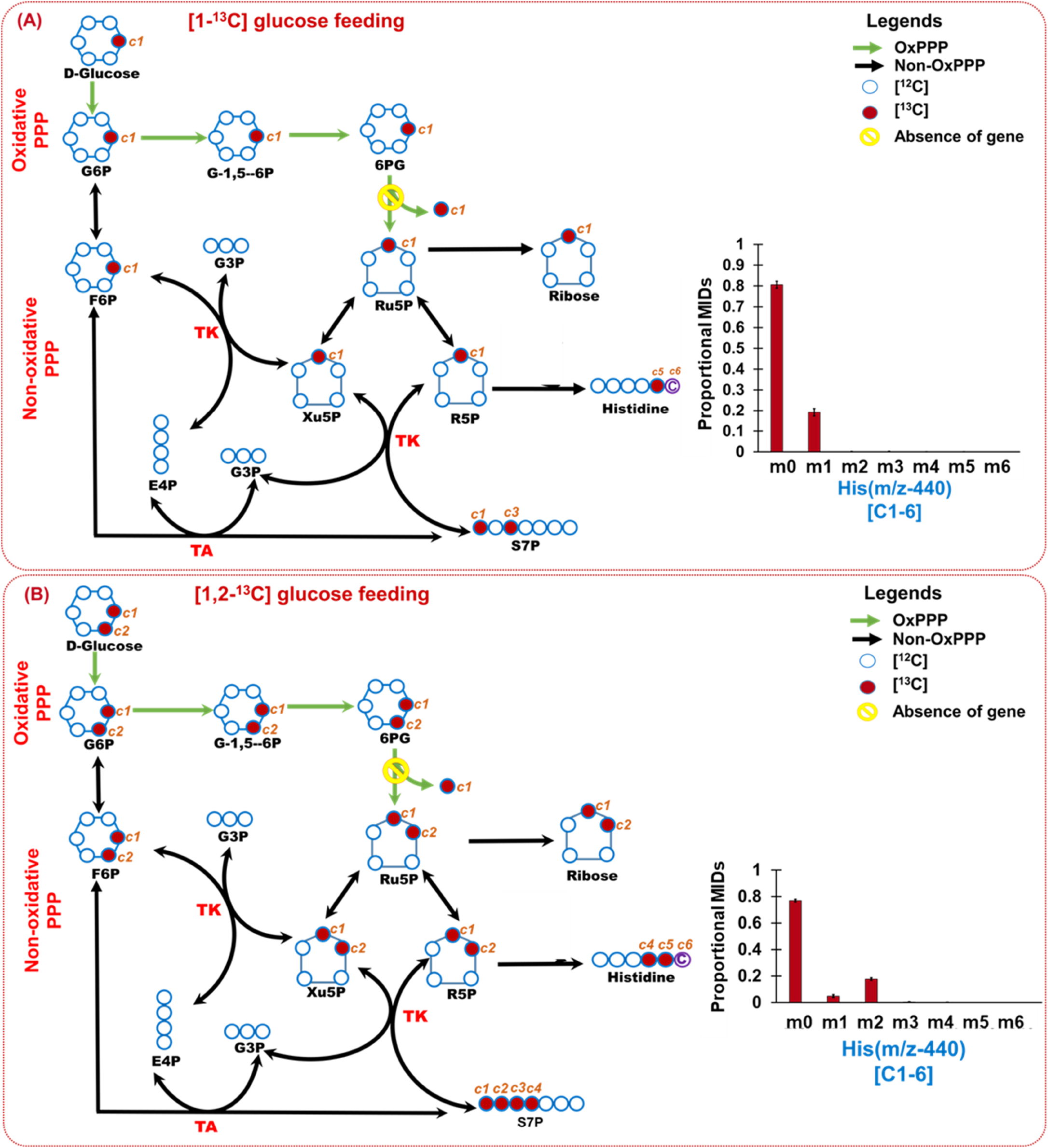
Movement of ^13^C label through OxPPP and Non-OxPPP in *R. solanacearum* F1C1. The carbon transition and label incorporation in the pathway intermediates were interpreted from the mass isotopomers of histidine derived from cells parallelly fed with [1-^13^C]glucose (A) and [1,2-^13^C]glucose (B). The mass isotopomer distribution in histidine m/z 440 (C 1-6) shows Non-OxPPP is the predominant route of glucose oxidation with no appreciable activity of OxPPP. [1- ^13^C]glucose oxidation via OxPPP would result in unlabelled histidine whereas NonOxPPP would result in one carbon labelled (m+1) histidine which is the case in these cells. Similarly, it is observed that cells fed with [1,2-^13^C]glucose exhibited higher levels of m+2 in histidine feasible via Non-OxPPP. See result section for details). Abbreviations: G-1,5-6P: Glucono-1-5-lactone-6phosphate; 6PG: phosphogluconate; Ru5P: Ribulose 5 phosphate; R5P: Ribose 5 phosphate; Xu5P: Xylulose 5 phosphate; F6P: Fructose 6 phosphate; G3P: Glyceraldehyde 3 phosphate; E4P: Erythrose 4 phosphate; S7P: Sedoheptulose 7 phosphate; PRPP: phosphor ribose diphosphate; MID: Mass isotopomer distribution

When cells were fed with [1-^13^C]glucose, it was observed that alanine (19.5% average ^13^C), valine (11.5%), glycine (3.5%) and histidine (4%) were labelled while no ^13^C enrichment was observed in serine, phenylalanine and tyrosine. Unlabelled serine, phenylalanine and tyrosine indicates that carbon movement is not via glycolysis. The higher labelling in alanine m/z 260 ([M-57] fragment ion contain C1-3 in the carbon backbone having 19.5% average ^13^C) in comparison to the alanine m/z 232 ([M-85] fragment ion with C2-3 in the carbon backbone with no significant label) indicates that the C1 position of alanine (or its precursor Pyruvate) is labelled, which is mainly possible via the ED pathway (Figure 3, Figure 4, Supplementary Table S3). The ED pathway activity as well as the lack of glycolytic activity was further supported by [1,2- ^13^C]glucose feeding data of Alanine m/z 260 (C1-3, 40% average ^13^C), Alanine m/z 232 (C2-3, 29% average ^13^C), Serine m/z 390 (C1-3, 2% average ^13^C) and Serine m/z 362 (C2-3, 1% average ^13^C) (Figure 3, Supplementary Table S3).

If glycolysis was active, we would have observed ^13^C incorporation in serine (or PGA, G3P) and C3 position of Alanine (or Pyruvate) from [1-^13^C]glucose and [1,2-^13^C]glucose feeding experiments, which was not the case (Figure 4A, 4B). We also checked the ^13^C enrichment in glycerol (m/z 377, mapped to glyceraldehyde 3-phosphate, nearest measurable product than serine) from [1-^13^C]glucose and [1,2-^13^C]glucose feeding experiments and found no significant labelling (Supplementary Figure S3) and hence further supporting inactive glycolysis. This clearly provides evidence that ED pathway bypasses glycolysis where in the labelled C-1 of glucose ends in pyruvate and subsequently in alanine and valine. The analysis clearly supports the notion that the lack of *pfk* gene in *R. solanacearum* renders inactivity of glycolysis with ED pathway as one of the preferred glucose oxidation routes.

We also observed that Non-OxPPP bypasses OxPPP as evidenced from both [1- ^13^C]glucose and [1,2-^13^C]glucose feeding experiments (Figure 5). The activities of PPP pathways were tracked retrobiosynthetically from histidine (m/z 440 (C1-6) whose carbons are derived from ribose5P and formate) and ribose (m/z 160 (C1-2)). Mass isotopomer distribution of histidine (Figure 5, Supplementary Table S4) and relative proportions of mass ion abundances of ribose (Supplementary Figure S3) has shed light on the PPP. From the cells fed with [1-^13^C]glucose, the labelling was observed in the mass isotopomer m+1 in histidine (m/z 440), which is possible by the activity of reductive pentose phosphate pathway (i.e Non-OxPPP). If OxPPP was active, it was expected that there would be loss of C1 from [1- ^13^C]glucose and we would obtain unlabelled ribose and histidine which is not the case. However, if both Non-OxPPP and OxPPP are active, we can still expect label incorporation in m+1 of histidine (m/z 440) and hence suggests that [1-^13^C]glucose feeding alone cannot confirm the inactivity of OxPPP. To further confirm the relative contributions of Non-OxPPP and OxPPP, the cells were also fed with [1,2-^13^C]glucose. The label incorporation showed higher levels of m+2 in comparison to m+1 which confirms that ^13^C label in ribose and histidine is mainly contributed via Non-OxPPP (Figure 5, Supplementary Figure S3). The data supports that Non-OxPPP is predominantly active in comparison to OxPPP possibly due to the lack of *gnd* gene (or appreciable OxPPP activity) in *R. solanacearum* strains as confirmed via comparative genome analysis (Figure 1).

## Discussion

*Ralstonia solanacearum* is a devastating phytopathogen and it is very important to understand its complete central metabolic network, mainly the elucidation of carbon metabolism. With the aim to investigate the intactness of all the central carbon pathways in *R. solanacearum* we undertook pathway comparisons of different strains (representing all the four phylotypes I,II, III and IV) derived from 53 genomes available in NCBI, Microscope/Genoscope and KEGG against the recently reported F1C1 strain (21), that we sequenced and annotated. In addition, 10 strains covering *Ralstonia* species other than *Ralstonia solanacearum* were considered for comparative pathway analysis (Supplementary Table S1).

It was observed that most of the *Ralstonia spp*. have some genes missing in glycolysis and OxPPP pathway raising the curiosity about how this genus sustains its metabolism. All the *R. solanacearum* strains lacked *pfk* gene that encodes an important regulatory enzyme phosphofructokinase-1 (EC 2.7.1.11, KO0850), which indicates possible bypass of glycolytic pathway by alternative routes. Also, *gnd* gene coding for phosphogluconate dehydrogenase enzymes (EC 1.1.1.44 and EC 1.1.1.343, KO0033) is absent in all *R. solanacearum* strains indicating incomplete OxPPP (Figure 1). *R. solanacearum* was earlier placed in the genus *Pseudomonas* (7) and it was reported that most of *Pseudomonas* species lack glycolysis (22). *Cupriavidus necator (previously classified as Ralstonia eutropha*) lack enzyme PFK-1 and follow ED pathway as an alternative to glycolysis (23). Although the possibility of alternative pathways seems evident from genome annotations, experimental validation of *in vivo* activities providing evidence of the absence of glycolysis and OxPPP in *R. solanacearum* is warranted. We selected ^13^C isotope labelling approach to get deeper insights of central carbon metabolic pathways and to validate the by-pass pathways as well as confirm the lack or extent of glycolytic and OxPPP activities in *R*. *solanacearum* F1C1. Parallel tracer feeding experiments using ^13^C Glucose (99%[1-^13^C]- or 99%[1,2-^13^C]- or 40%[^13^C_6_]-glucose) were undertook for comprehensive insights into pathway activities. In terms of the approaches used, ^13^C incorporation in the carbons of amino acids retrobiosynthetically report on the ^13^C incorporations in precursors and other central metabolites. Parallel ^13^C labelling experiments (using [1-^13^C]-, [1,2-^13^C]-, [^13^C_6_]-glucose) were conducted which is standard practice and is robust.

Cells fed with [1-^13^C] glucose has ^13^C enrichment in pyruvate but there is no ^13^C incorporation in serine and glycerol (Figure 3, Figure 4, Supplementary Table S3 & S4, Supplementary Figure S3) indicating the lack of glycolysis and active ED pathway which is further supported by labelling in the C1 of alanine (19, 24). The equal distribution of average ^13^C (in %) in serine and alanine from [^13^C_6_] glucose feeding is another indication of ED pathway activity mainly (Figure 3, Supplementary Table S3 & S4).

Higher ^13^C enrichment in [M-57] fragment of alanine m/z (260) compared to [M-85] fragment (alanine m/z (232)), when [1-^13^C] glucose was used, is the confirmation of active ED pathway (25). Similar results were reported in *Dinoroseobacter shibae* fed with [1-^13^C] glucose (25).

The activity of Oxidative pentose phosphate pathway is questionable in *R.solanacearum* as comparative genome analysis highlighted the absence of key gene 6-phosphogluconate dehydrogenase (*gnd*, EC 1.1.1.44) (Figure 1). To validate this we undertook parallel [1-^13^C]- and [1,2-^13^C] glucose feeding and measured the ^13^C incorporation in histidine and ribose, that can retro-biosynthetically report on the PPP activities. The carbon precursors for histidine biosynthesis are, 5-phospho-ribosyl-a-pyrophosphate (PRPP) and the formyl group of N10-formyl-tetrahydrofolate (THF) (24). In OxPPP, the first (labelled) carbon in [1-^13^C]glucose is generally released as ^13^CO_2_ during the conversion of 6-phosphogluconate into D-ribulose-5-phosphate by *gnd*. In *R.solanacearum*, if OxPPP is predominantly active than it is expected to not get any ^13^C in histidine. However, ^13^C incorporations in both histidine and ribose were observed which explains that the predominant activity is due to the Non-oxidative or reductive PPP (Non-OxPPP). (Figure 5, Supplementary Table S4, Supplementary Figure S3). To further confirm that ^13^C label is from Non-OxPPP, bacterial cells were also fed with [1,2- ^13^C] glucose. It was observed that the mass isotopomer abundances in histidine (m+2) and relative proportions of mass representing ribose (m+2) were higher than m+1 respectively (Figure 5, Supplementary Figure S3) which confirms that Non-OxPPP is predominantly active than the OxPPP possibly due to the lack of *gnd* gene in *R.solanacearum*.

In addition, on close examination of the publicly available transcriptome data of *R. solanacearum* strains GMI1000 and UW551 derived from in-planta and rich media growth (4), it was found that pfk-1 and gnd gene expressions are not reported (while several other genes such as that of ED pathway are expressed) potentially owing to their absence. Similarly, the Transcriptome analysis by Puigvert et al 2017 has no reports of *gnd* and *pfk-1* genes in homologous gene analysis of *R. solanacearum* strains whereas ED pathway genes were reported (26). These further supported our findings that *gnd* and *pfk-1* genes are inactive or altogether absent in *R solanacearum* and that mainly glucose oxidation bypasses glycolysis and OxPPP in favour of ED-Pathway and Non-OxPPP.

Overall, this work supported by comparative pathway analysis from genomes, parallel ^13^C labelling analysis suggests that ED pathway and Non-OxPPP are the main metabolic routes followed by *R. solanacearum* for glucose oxidation. ED pathway plays significant roles in the activation of virulence factors and inhibition of biofilm formation (27). Biofilm aids in survival of the pathogen and help to enter in the host but inside the host, bacterial cells disperse and form individual colonies for infection. Although ED pathway produced one ATP (less than glycolysis), there are several microbes that prefer ED pathway over glycolysis probably due to some evolutionary advantage. To explore the reason, Flamholza and coworkers (28) studied the energy yield and protein cost in glycolytic and ED pathways in prokaryotes. They hypothesized that the enzymatic synthesis cost also plays a key role in the choice of the pathway used for glucose catabolism. The use of the ED pathway results in a lower protein synthesis cost to catabolize the same amount of glucose than glycolysis (28). Similarly, the lack of OxPPP activity would starve the cells of NADPH levels which may be compensated by alternative pathways producing NADPH or reducing the cellular needs of this valuable resource. It remains to be seen how *R. solanacearum* adjusts to its metabolic demands under varied nutritional niches it encounters. The findings reported here will be on immense relevance while defining the metabolic phenotypes of *R. solanacearum*. This is the first comprehensive ^13^C labelling study in *R. solanacearum* with focus on deciphering its glucose oxidation and mapping of central carbon metabolic pathways in *R. solanacearum*.

## Materials and Methods

### Chemicals

The labelling substrates, [1-^13^C]glucose (99 atom %), [1,2-^13^C]glucose (99 atom %), and [^13^C_6_]glucose (99 atom %) were purchased from Sigma-Aldrich. Derivatization agents, TBDMS (N-methyl-N-(t-butyldimethylsilyl) trifluoroacetamide + 1% t-butyl-dimethylchlorosilane) and MSTFA (N-methyl-N-(trimethylsilyl) trifluroacetamide) were also purchased from Sigma-Aldrich. Media components and other reagents were purchased from Himedia and Sigma-Aldrich.

### Softwares

KEGG mapper, Agilent Chemstation, MassHunter, IsoCorr, Metalign, Amdis (NIST). Online resources were also used.

### Analytical equipments

GC-MS (Agilent), Microplate reader (ELx 808, Bio-tek instruments Inc).

## Experimental procedure

### Comparative metabolic pathway analysis

For comparative analysis of metabolic pathways, genome sequence of 53 strains of *Ralstonia solanacearum* including F1C1(unpublished data), which belong to different phylotypes (represent all four phylotypes, I II III and IV) were selected (Supplementary Table S1). Six other *Ralstonia* spp. (covering 10 strains) namely, *R. syzygii* strain-R24, *R. insidiosa* strain-FC1138, *R. eutropha* (now renamed as *Cupriavidus necator* strain-H16, strain JMP134), Banana blood disease strain R229, *R. pseudosolanacearum strain RS 476, R. mannitolilytica strain SN82F48* and *R. picketti* strain-12D, strain-DTP0602, strain-12J; along with *E. coli strain DH10B* (as positive control) were selected. Central metabolic pathways were compared based on KEGG pathway mapping (15). For the organisms not present in the KEGG database, the multifasta amino acid sequences were downloaded from Genoscope/Microscope (16) or NCBI (29). Then KO identifiers from these amino acid fasta files were generated using BlastKOALA (BLAST-based KO annotation and KEGG mapping) (30). Comparative pathway analysis and visualisation of selected *Ralstonia spp*. along with *E. coli* was done using KEGG pathway reconstruct tool (30)

### Cell growth and maintenance

*R. solanacearum* F1C1 culture was maintained in BG medium in 50 ml falcon tubes at 28°C on an orbital shaker running at 180 rpm (31, 32). For pathway mapping experiment, cells were grown in BG medium for 24 hrs untill the OD600 reaches to 1. Cells were pelleted down by using centrifuge at 5000 rpm for 5 mins. Cells were transferred to minimal media and grown for 24hrs to adapt in minimal media. Cells were further harvested, washed with sterile water and redistributed in minimal medium supplemented with labelled Glucose (0.5% w/v) four replicates each. The composition of minimal medium (14) is as follows (gL^−1^): FeSO_4_-7H_2_O, 1.25×10^−4^; (NH_4_)2SO_4_, 0.5; MgSO_4_-7H_2_O,0.05; KH_2_PO_4_, 3.4. The pH was adjusted to 7.0 with 1M KOH. Cells were grown in minimal media with different combinations of labelled isotopes ([1-^13^C]-, [1,2-^13^C]-, 40%[^13^C_6_]-, [12C]-Glucose; four technical replicates each, Supplementary Figure1). Cells were harvested at 18 h during mid-exponential phase that represent pseudo-steady state condition (Supplementary Figure S2).

### Acid hydrolysis and Derivatization of amino acids

Cell pellets (2 mg each) were acid hydrolysed by suspending them in 500 μL of 6M HCl for 18 hr at 100 °C to release the amino acids (33). 50 μL acid hydrolysate was then dried in speed-vacuum (Thermo-scientific,Waltham, MA, USA) to ensure complete removal of water. The amino acids extracts were derivatised by TBDMS (34). To obtain the TBDMS derivatives, the vacuum dried acid hydrolysates of amino acids were first dissolved in 30 μl of pyridine (Sigma Aldrich) and incubated at 37 °C, shaking at 900 rpm for 30 min. Then 50 μl of MtBSTFA + 1% t-BDMCS (N-methyl-N-(t-butyldimethylsilyl) trifluoroacetamide + 1% t-butyl-dimethylchlorosilane, Regis Technologies Inc) was added and incubated the samples at 60 °C with shaking at 900 rpm for 30 min on a thermomixer. The derivatised samples were centrifuged for 10 min at 13000 rpm to pellet down any insoluble material and transferred the supernatant to glass vials for GC-MS and sealed with a septum cap.

### Sample preparation for ribose labelling

^13^C enrichment in ribose was measured as suggested by Long and Antoniewicz (35). The dry cells (2 mg) were acid hydrolysed with 50 μl of 6N HCL for 30 mins and then it was diluted to 1N by adding 250 μl of distilled water and incubated for 1 hr in fume hood. The reaction was neutralized by adding 40 μl of 5N NaOH. The reaction mixture was centrifuged at 14000 rpm for 10 mins. 50 μl of supernatant was dried in speed-vacuum (Thermo-scientific). To obtain the TMS derivatives, the vacuum dried extracts were dissolved in 35 μl of freshly prepared methoxyamine hydrochloride (MeOX, Sigma Aldrich) in pyridine (20 mg/ml) and incubated on a thermoshaker at 37 °C, 900 rpm for 2 hrs. After that 49 μl of MSTFA was added and incubated the samples at 37 °C with, 900 rpm for 30 min on a thermomixer (36). The derivatised samples were centrifuged for 10 min at 13000 rpm and supernatant was transferred to glass vials for GC-MS.

### Gas chromatography-mass spectrometry

The GC-MS measurements were performed on an Agilent 7890B GC, electron impact ionisation (70 eV) equipped with an Agilent HP 5-ms ultra-inert column (Agilent 19091S-433UI, 30m x 250μm x 0.25 μm dimensions) at the facility in BioX center at IIT Mandi. In GC-MS, 1 μl sample volume was taken for injection. For TBDMS derivatised amino acid hydrolysate, the initial oven temperature was constant at 120 °C for 5 min, then a 4 °C /min ramped to 270 °C, held for 3 min, then a 20 °C/min ramped to 320 °C and held for 1 min. The carrier gas (Helium) flow was maintained at 1.3 ml min^−1^. The spectra were recorded with a scanning range of 30 to 600 mz^−^1 for a total run time of 49 mins. For MeOX-TMS derivatised samples, oven temperature was constant at 50 °C for 5 mins and then ramped to 200 °C with 10 °C /min and held for 10 min. Temperature was again ramped to 300 °C with 5 °C /min and held for 10 min. After that temperature was decreased to 70 °C with 100 °C /min. The carrier gas (Helium) flow was 1.3 ml min^−1^. MassHunter (Agilent Technologies, USA) was used to control the data acquisition parameters (both GC separation and mass spectrometry) during all the sample runs.

### Metabolite identification and mass isotopomer data handling

The raw GC-MS spectra need to be baseline corrected at first for accurate assessment of mass isotopomer distributions in metabolites. The raw files from GC-MS was baseline corrected using MetAlign software (37) with its default parameters. Metabolite identification was done using NIST (National Institute of Standards and Technology, Maryland). The intensity of the mass ions of each amino acid fragment (38) was obtained by using Agilent chemstation software. The MIDs of each fragment ion obtained from averaged mass spectra were corrected for the presence of naturally occurring heavy isotopes attached to the carbon backbone of the derivative using mass correction software IsoCor (39).The corrected MID values were subjected to calculate the average ^13^C abundance in each fragment (34)

### Statistical Analysis

Four replicates were used in all the analysis and data was present as the mean ± SD for each experiment. The statistical significances were performed using Student’s t test.

## Acknowledgements

All authors acknowledge the financial support from Department of Biotechnology (DBT), Govt. of India provided to Tezpur University and IIT Mandi (Twinning grant - BT/PR16361/NER/95/192/2015; IITM/DBT/SKM/126). SKM and TPS also acknowledges the seed grant provided by IIT Mandi (ITM/SG/SKM/48). PJ thanks CSIR-UGC JRF fellowship for supporting her doctoral studies.

## Author Contributions

PJ and SKM conceived and designed the experiments; PJ performed the experiments. PJ, CJ and TP contributed in comparative genome analysis. SS, SR provided strain, its genome sequence and contributed in annotation along with PJ and SKM. PJ, MS, and SKM analysed ^13^C data and wrote the paper.

## Supplementary Materials

**Figure S1** Experimental workflow adopted for central metabolic pathway mapping by isotopic tracer feeding and GC-MS data analysis towards average ^13^C label incorporation in proteinogenic amino acids

**Figure S2** The growth curve of *Ralstonia solanacearum* cells in minimal media containing glucose (0.5% w/v) as carbon source.

**Figure S3** The relative proportion of Ribose fragments (m/z-160) and Glycerol fragments (m/z-377) of *Ralstonia solanacearum* F1C1 subjected to minimal media supplemented with [^12^C_6_]-, [1-^13^C]- and [1,2-^13^C]-glucose.

**Table S1** Comparative pathway analysis of *Ralstonia spp*. highlighting the absence of key pathway genes.

**Table S2** Amino acid fragments of *R. solanacearum* with their respective Peak no, elution time, derivative and specific ions (m/z) obtained from GC-MS

**Table S3** Average ^13^C abundance (in %) of amino acid fragments of *Ralstonia solanacearum* F1C1 subjected to minimal media supplemented with [1-^13^C]-, [1,2-^13^C]- and [^13^C_6_] glucose.

**Table S4** Mass isotopomer distributions (MID) of amino acid fragments obtained from *Ralstonia solanacearum* F1C1 subjected to minimal media supplemented with [1-^13^C]-, [1,2-^13^C]- and [^13^C_6_] glucose.

---

**Supplementary Figure S1.** Experimental workflow adopted for central metabolic pathway mapping by isotopic tracer feeding and GC-MS data analysis towards average 13C label incorporation in proteinogenic amino acids *R. solanacearum* cells were grown overnight in BG media till optical density (OD_600_) reaches to 1. Cells were harvested by centrifugation at 5000 rpm and then transferred to minimal media to adapt for 24hrs, again harvested and redistributed in minimal media with Isotopic tracers i.e. 40%[^13^C_6_]glucose, [1-^13^C_6_]glucose and [1,2-^13^C_6_]glucose along with ^12^C substrates as control (four replicates each). Cells were harvested after 18hr of inoculation and were acid hydrolysed using 6M HCl. Protein hydrolysates were derivatised by TBDMS (Tertiary butyl dimethyl silane) derivatization for GC-MS analysis. The TIC (Total Ion Chromatograms) of all amino acids were corrected for natural isotope correction and validation was based on average ^13^C in unlabelled fragments. Validated amino acid fragments were then used to map the central metabolic pathways.

**Supplementary Figure S2.** The growth curve of *Ralstonia solanacearum* cells in minimal media containing 0.5% (w/v) glucose as carbon source. For metabolic pathway mapping cells were harvested at 18 h during mid-exponential phase that represent pseudo-steady state condition.

**Supplementary Figure S3.** The relative proportion of mass ions from fragments of Ribose (m/z-160) (3a and 3b) and Glycerol (m/z-377) (3c and 3d) of *Ralstonia solanacearum* F1C1 subjected to minimal media supplemented with either [^12^C_6_], [1^13^C]- or [1,2-^13^C]-glucose. The mean relative abundance of mass ions and corresponding standard deviations from 4 replicates is presented. The significant difference in relative proportions were analysed by Student’s t test * P < 0.5, ** P < 0.01, *** P < 0.001, and **** P < 0.0001 in ribose (Figure 3a and 3b) and * P < 1, ** P < 0.5, *** P < 0.001, and **** P < 0.0001 in Glycerol (Figure 3c and 3d). The mass fragments are not corrected for ^13^C natural abundances.

**Supplementary Table S1.** Comparative pathway analysis of 53 *Ralstonia solanacearum* strains and 10 strains from different *Ralstonia spp*. along with *E. coli* based on presence and absence of key pathway genes for glucose oxidation namely, glycolysis, ED-pathway and Pentose Phosphate Pathway. Black colored boxes represent the absence of genes. All *Ralstonia spp*. strains studied have absence of *pfk-1* gene (support glycolysis) except for *R. picketti* while *gnd gene* (support OxPPP) is absent in all the *Ralstonia spp. E. coli* was used as control which has all the glucose oxidation *pathways (*glycolysis, ED-pathway and Pentose Phosphate Pathway) intact.

**Supplementary Table S2.** Amino acid fragments of *R. solanacearum*. All the detected 15 amino acids with their respective Peak no, elution time (Figure 1), derivative and specific ions (m/z) obtained from GC-MS are tabulated

**Supplementary Table S3.** Average 13C abundance (in%) of *Ralstonia solanacearum* F1C1 strain (valid, [M-57] & [M-85] amino acids fragment ions and standard deviation from 4 replicates) subjected to minimal media supplemented with [^12^C_6_]-, [^1-13^C]-, [^1,2-13^C]- and [^13^C_6_]-glucose.

**Supplementary Table S4.** The Mass Isotopomer Distributions (MID) of amino acid fragments (valid, [M-57] and/or [M-85] and standard deviation from 4 replicates) of *Ralstonia solanacearum* F1C1 subjected to minimal media supplemented with [^12^C_6_]-, [^113^C]-, [^1,2-13^C]- and [^13^C_6_]-glucose

